# Obstacles to the reuse of study metadata in ClinicalTrials.gov

**DOI:** 10.1101/850578

**Authors:** Laura Miron, Rafael S. Gonçalves, Mark A. Musen

## Abstract

Metadata that are structured using principled schemas and that use terms from ontologies are essential to making biomedical data findable and reusable for downstream analyses. The largest source of metadata that describes the experimental protocol, funding, and scientific leadership of clinical studies is ClinicalTrials.gov. We evaluated whether values in 302,091 trial records adhere to expected data types and use terms from biomedical ontologies, whether records contain fields required by government regulations, and whether structured elements could replace free-text elements. Contact information, outcome measures, and study design are frequently missing or underspecified. Important fields for search, such as c*ondition* and *intervention*, are not restricted to ontologies, and almost half of the *conditions* are not denoted by MeSH terms, as recommended. Eligibility criteria are stored as semi-structured free text. Enforcing the presence of all required elements, requiring values for certain fields to be drawn from ontologies, and creating a structured *eligibility criteria* element would improve the reusability of data from ClinicalTrials.gov in systematic reviews, metanalyses, and matching of eligible patients to trials.

## Introduction

Over the past two decades, the scientific community has increasingly recognized the need to make the protocols and results of experiments publicly accessible so that data can be reused and analyzed. However, the findability and meaningful reuse of data are often hampered by the lack of standardized metadata that describe the data. Biomedical metadata are typically records of key–value pairs that are created when investigators submit data to a repository such as the Gene Expression Omnibus (GEO), Protein Data Bank (PDB), or NCBI BioSample repository. Metadata describe the source of the data (e.g., investigators, sponsoring organizations, data submission and update dates), the structure of datasets, experimental protocols, identifying and summarizing information, and other domain-specific information.

Clinical-trial registries are repositories of structured records of key–value pairs (“registrations”) summarizing a trial’s start and end dates, eligibility criteria, interventions prescribed, study design, names of sponsors and investigators, and prespecified outcome measures, among other details. As opposed to standards that enable researchers to share structural and administrative metadata about the format and provenance of datasets from clinical trials (e.g., the Clinical Data Interchange Standard Consortium’s (CDISC’s) Define.XML standard), registrations represent a class of descriptive metadata about the high-level characteristics of a clinical trial. The largest such registry is ClinicalTrials.gov,^1^ a Web-based resource created and maintained by the U.S. National Library of Medicine (NLM). The official purpose of ClinicalTrials.gov is to implement the requirements from a series of acts by the US Food and Drug Administration (FDA; Food and Drug Administration Modernization Act of 1997, Food and Drug Administration Amendments Act of 2007, FDAAA801 Final Rule) which mandated the registration of trials of controlled drugs and devices as a means to safeguard human subjects, but these registrations are also the largest collection of metadata about clinical trials in the world, and are increasingly being reused for other important purposes.

Clinical-trial registries such as ClinicalTrials.gov have become a crucial source of information for systematic reviews and other metanalyses. Recent studies recommend that systematic reviews include a search of clinical trial registries to identify relevant trials that are ongoing or unpublished^2–5^. Baudard et al.^6^ selected 116 systematic reviews that did not report a search of clinical trial registries and were able to find 122 additional randomized controlled trials in registries that increased the number of patients for 41 reviews and changed summary statistics by greater than 10% for 7 reviews. Hart et al.^7^ found that, for 41 drug trials, the inclusion of unpublished trial outcome data caused an increase in estimated efficacy in 19 trials (median change 13%) and a decrease in estimated efficacy in 19 trials (median change 11%). Including registry metadata in systematic reviews can help to identify selective reporting bias by comparing published outcomes to prespecified outcomes^8,9^, and adverse events are more likely to be reported in clinical trial registries than in published literature^10,11^. Records from ClinicalTrials.gov are also commonly used in metanalyses about trends in sources of funding for trials^12–16^, diseases and interventions studied^13,17–21^, study design^17,19,22^, time to publication following study completion^11,23–26^, geographical availability of trials sites^27–32^, and the causes of delays and early terminations in studies^33–40^.

ClinicalTrials.gov records, like metadata records from other widely used biomedical data repositories,^41,42^ are plagued by quality issues. Several studies have analyzed ClinicalTrials.gov records for missing fields required by the Food and Drug Administration Amendments Act of 2007, which governs US trial registries, and the World Health Organization (WHO) minimum data set, which provides guidelines for registries internationally^43–46^. Chaturvedi et al.^47^ found that information about the principal investigators of trials in ClinicalTrials.gov are inconsistent both within multiple occurrences in the same record and across records. Tse et al.^48^ identify additional obstacles to clinical trial data reuse: follow-up studies are not always linked to the original study, records can be modified by the responsible party at any time, standards include both mandatory and optional data elements, and the presence of records in the database is biased by reporting incentives.

While these quality issues with ClinicalTrials.gov records are known, we have found no analyses of whether trial records have structural characteristics that impede the reuse of the metadata, akin to the problems found in other biomedical repositories. Hu et al. examined the quality of the metadata that accompany data records in the Gene Expression Omnibus (GEO) tnd found that they suffered from type inconsistency (e.g., numerical fields populated with non-numerical values), incompleteness (required fields not filled in), and the use of many syntactic variants for the same field (e.g., “age, Age, Age years, age year”)^41^. In past work, we documented similar quality issues with the NCBI BioSample and EBI BioSamples repositories: records used many syntactic variants for the same field, and values did not conform to the expected type, including where ontology term identifiers are expected^42^. Both of these studies also concluded that the metadata entry pipelines, which allowed the submission of user-defined fields and provided limited automated validation, contributed significantly to the quality of records.

In this analysis, we investigated whether clinical-trial metadata values conform to expected data types, whether values are ontology terms where recommended, and whether unstructured free-text elements could be replaced with structured elements. We also determined updated counts of records missing required elements. Finally, we examined the data-entry pipeline for ClinicalTrials.gov, mediated by software known as the Protocol Registration System (PRS)^49^. Since the PRS does not allow user-defined fields, and all ClinicalTrials.gov records conform to a single XML schema, we do not analyze the use of variants for field names.

We found that automated validation rules within the PRS have been successful at enforcing type restrictions on numeric, date, and Boolean fields, and fields with enumerated values. However, fields for entry of contact information, principal investigators, study design, and outcome measures are frequently missing or underspecified. Values for fields commonly used in search queries, such as *condition*(s) and *intervention*(s), are not restricted to ontology terms, impeding search. Eligibility criteria, which could be used to facilitate the matching of patients to applicable clinical trials if stored in a structured format, are currently stored as semi-structured free-text and cannot be used to query the repository.

## Background

ClinicalTrials.gov was first released by the NLM in 2000, and it consists of a clinical-trial registration repository, the PRS for submitting records, and a user-facing website. The search portal of the user-facing website allows queries based on conditions, interventions, sponsors, locations, and other fields within the metadata.

### Data Entry: Protocol Registration System

The data-entry system for ClinicalTrials.gov is PRS, a Web-based tool that provides form-based entry pages and that also includes a quality-review pipeline with both automated validation rules and support for manual review by a member of the NLM staff. The PRS form-based entry system employs several methods to improve data quality. Markers by each field name indicate whether the element is required. Radio buttons are used for entry of Boolean values and drop-down menus are associated with fields that have enumerated values (**Figure 1**). Automated validation messages of four possible warning levels (**Table 1**) appear when errors or potentially wrong values are detected. The author may only submit the record for manual review when all errors are resolved.

**Figure 1.**
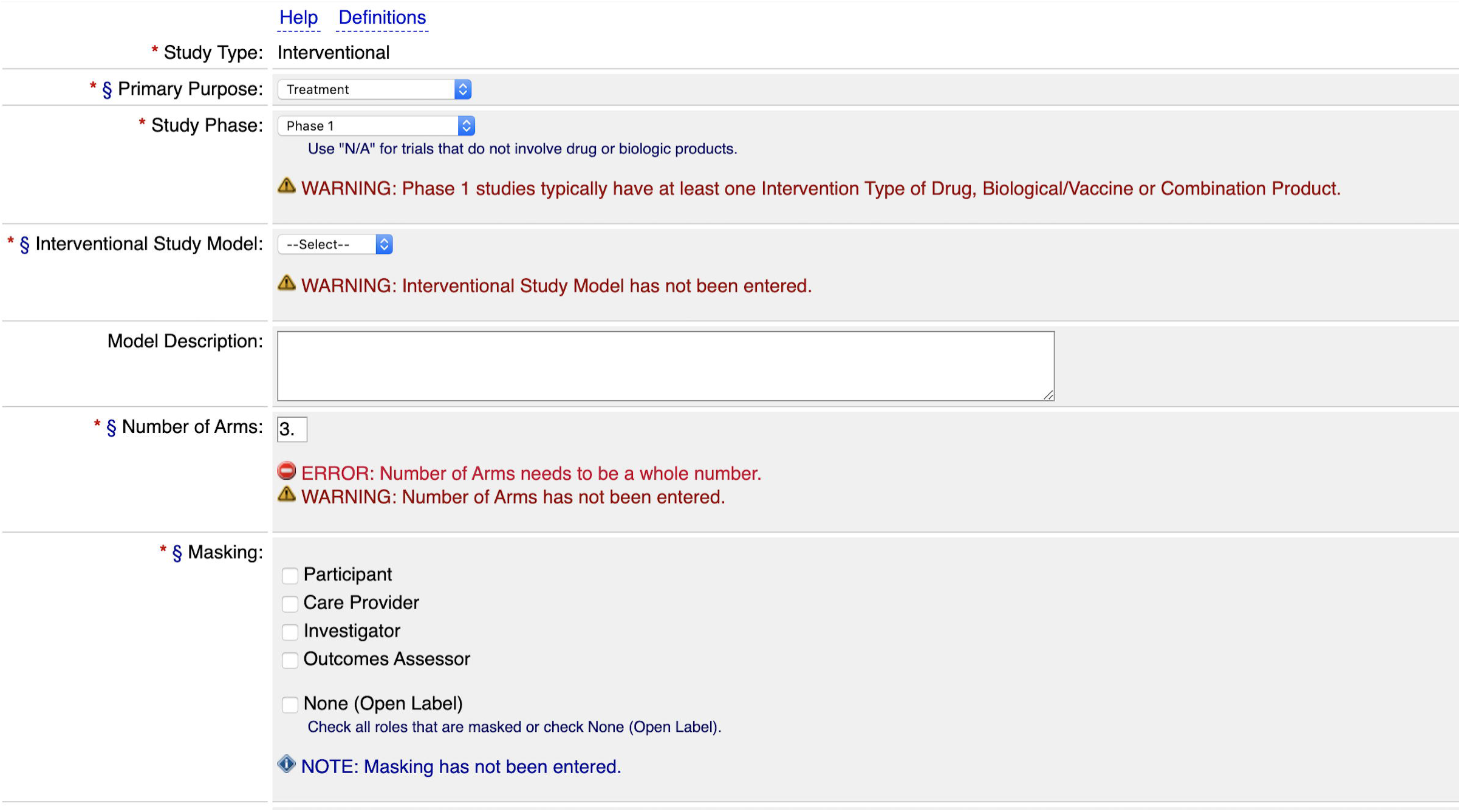
Data entry form in the PRS system. One of several form pages for entering data in the PRS. Red asterisks (**\***) indicate required fields; red asterisks with a section sign (**\***§) indicate fields required since January 18, 2017. Additional instructions are provided for ‘study phase’ and ‘masking’ fields, and automated validation messages of levels ‘Note’, ‘Warning’ and ‘Error’ can be seen. A validation rule ensures that the value for ‘number of arms’ is an integer. Another rule checks both the chosen value for ‘study phase’ (Phase 1), and the (lack of) interventions that are enumerated on a separate page of the entry system. However, there is unexplained inconsistency in the warning levels for missing required elements (missing ‘masking’ generates a ‘note’, while missing ‘interventional study model’ and ‘number of arms’ both generate ‘warnings’).

**Table 1.** Warning Levels in the Protocol Registration System.

| Type | Explanation |
| --- | --- |
| Error | Problems that must be addressed (e.g., missing required content, internal inconsistency) |
| Warning | Items that are FDAAA <u>required</u> or FDAAA <u>may be required</u> (e.g., Study Start Date data element) |
| Alert | Problems that need to be addressed |
| Note | <u>Potential</u> problems that should be reviewed and corrected, if possible (if not possible, then may ignore) |

### Characteristics of High-Quality Metadata

Communities of investigators in several biomedical domains have defined a “minimum information standard”, or list of required fields, for metadata about a particular type of experiment (e.g., the Minimum Information About a Microarray Experiment (MIAME)^50^). FAIRsharing.org^51^ provides a registry of such standards for metadata in various scientific disciplines. High quality, “complete” metadata contain values for all fields required by the relevant minimum information standard. For clinical-trial data, two main policies govern minimum information standards:

1. International Committee of Medical Journal Editors (ICMJE)/ World Health Organization (WHO) trial registration dataset^52^
2. Section 801 of the Food and Drug Administration (FDA) Amendments Act of 2007 (FDAAA801)^53^, and its Final Rule (42 CFR Part 11)^54^, which updated and finalized required element definitions

In principle, the WHO trial registration dataset^52^ applies to all interventional trials in the world, whereas FDAAA801 applies only to interventional trials of controlled drugs and devices within the United States. In practice, however, ClinicalTrials.gov is the largest international repository of both types of trials, and its required element definitions correspond to those defined in FDAAA801. The ICMJE and WHO therefore accept trials that are fully registered in ClinicalTrials.gov as meeting their standard, and release an official mapping of WHO data elements to ClinicalTrials.gov data elements^55^.

Metadata schemas often require that values for certain fields be drawn from a particular biomedical ontology in order to prevent the usage of synonyms and to provide a defined range of values that can be used to query the metadata. Use of terms from well-known domain-specific ontologies is one of the fundamental guidelines enumerated by the FAIR principles for making scientific data and metadata Findable, Accessible, Interoperable, and Reusable^56^.

Where appropriate, values should be defined using globally unique and persistent identifiers, such as the URIs of terms in (a particular version of) an ontology (e.g., http://purl.bioontology.org/ontology/MESH/D003920). A globally unique identifier denotes a term unambiguously, regardless of homonyms, and a persistent identifier gives researchers who consume metadata a reliable pointer to information about the term, such as labels, synonyms, and definitions.

## Methods

ClinicalTrials.gov records are available as Web pages—accessible through the system’s search portal—and as XML files (https://ClinicalTrials.gov/AllPublicXML.zip). We downloaded all public XML trial records (n=302,091) on April 3, 2019. We conducted our analysis of the PRS system using a test environment, which allows records to be created but never submitted, maintained by Stanford University. We conducted several analyses on the XML records: We enumerated all fields that expect values to conform to a simple type (integer, Boolean, date, or enumerated values) or to be drawn from an ontology, and tested whether values adhere to type expectations. To evaluate completeness, we counted the numbers of records missing all fields required by FDAAA801. We used regular expressions to test whether values for *eligibility criteria* conformed to the expected semi-structured format.

ClinicalTrials.gov contains three kinds of records: those for *interventional* trials (subjects are prospectively assigned interventions), for *observational* trials (outcomes are retrospectively or prospectively observed, but interventions are not prescribed; may additionally be designated *patient registries*), and *expanded-access* records. Expanded-access records exist in conjunction with existing records for interventional studies, in cases where study sponsors also administer the experimental interventions to patients who are ineligible for the main cohort. Because FDAAA801 only defines required fields for interventional trials, we conducted analyses of missing fields only on the set of 239,274 interventional records, and conducted all other analyses on the full set of 302,091 records.

### Data Element Definitions and Schema

ClinicalTrials.gov data elements are defined in a free-text data dictionary and in an XML schema declaration (XSD). There is a separate data dictionary specification for interventional and observational records (https://prsinfo.ClinicalTrials.gov/definitions.html) and for expanded access records (https://prsinfo.clinicaltrials.gov/expanded_access_definitions.html), but most elements are common to all three types of records and they all conform to the same XSD. A table containing the exact mapping between element names in the data dictionary, XML element names, field names in FDAAA801, and WHO data element names is provided in the supplementary material. The mapping between data dictionary elements and XML elements is mostly trivial, but several dictionary/XML elements can map to a single field required by FDAAA801, and others are optional and not included in FDAAA801.

For our analyses of type consistency and usage of ontology terms, we considered all 149 elements listed in the two data dictionaries. For our analysis of completeness, we considered only 78 elements corresponding to the 41 required FDA elements. Unless otherwise specified, we refer to elements by the name provided in the data dictionary. Definitions for significant fields in our analyses are given in **Table 2**.

**Table 2.** Significant Fields in ClinicalTrials.gov. Definitions are adapted from the field definitions provided in the ClinicalTrials.gov data dictionary

|  |  |
| --- | --- |
| *Condition* | The name(s) of the disease(s) or condition(s) studied in the clinical study |
| *Intervention* | The intervention(s) associated with each arm or group, most commonly a drug, device, or procedure |
| *Eligibility Criteria* | A limited list of criteria for selection of participants in the clinical study, provided in terms of inclusion and exclusion criteria |
| *Outcome Measure* | A prespecified measurement used to determine the effect of experimental variables on human subjects in a clinical study |
| *Responsible Party* | Either a “Sponsor”, “Principal Investigator”, or “Sponsor-Investigator”; the party responsible for submitting information about a trial |
| *Overall Official* | Person responsible for the overall scientific leadership of the protocol, including study principal investigator |

### Adherence to Simple Type Expectations

We assigned each metadata field in the data dictionary to a category, and, for each category, we determined the type of validation that we would perform:

- Simple type (date, integer, age, Boolean) – Validate records against the XSD.
- Enumerated-values field (data dictionary provides enumerated list of acceptable values) – Programmatically check values against expected values from data dictionary.
- Ontology-controlled field – Validate values against the expected ontologies.
- Free text – Validation of *eligibility criteria* element only, discussed below

### Usage of Ontology Terms

We used the National Center for Biomedical Ontology (NCBO) BioPortal API^57^ to find ontology terms whose preferred names are exact matches for values for the *condition* and *intervention* fields. BioPortal is the most comprehensive repository of biomedical ontologies, and has a publicly accessible API that can be queried for data about ontology identifiers (e.g., unique identifier, source ontology) corresponding to the search term^57^. The *condition* and *intervention* fields within ClinicalTrials.gov records share characteristics of fields that could support and be improved by ontology restrictions on the allowed values: expected values for these fields are already likely to be found in well-known ontologies such as MeSH or RXNORM, unrestricted values for these fields are likely to introduce synonyms (e.g., the proprietary name and generic name for a drug), and they are important fields for querying the repository.

Currently, only the *condition* field is ontology-restricted in ClinicalTrials.gov. The data dictionary says to “use, if available, appropriate descriptors from NLM’s Medical Subject Headings (MeSH)-controlled vocabulary or terms from another vocabulary, such as the Systemized Nomenclature of Medicine–Clinical Terms (SNOMED-CT), that has been mapped to MeSH within the Unified Medical Language System (UMLS) Metathesaurus.”^58^ To test adherence to this restriction, we used BioPortal to search for exact matches for each term, restricted to the 72 ontologies in the 2019 version of UMLS. To evaluate the degree to which *intervention* field values were ontology-restricted, we queried all ontologies in BioPortal for exact matches for each intervention listed.

### Missing Fields

We counted the number of records with missing fields for 28 of the 41 fields required by FDAAA801, the statute that defines the minimum required elements for clinical trial registrations in the US. We ignored five fields that are conditionally required based on information unavailable to us (e.g., secondary outcome measures must be listed, but only if they exist) (*Pediatric Postmarket Surveillance, Other Names for Interventions, Post Prior to FDA Approval/Clearance, Product Manufactured in or Exported from the U*.*S*., *Secondary Outcome Measure Information*), three fields stored internally by ClinicalTrials.gov but not made public *FDA IND or IDE, Human Subject Protection Board Review Status, Responsible Party Contact Information*), three fields which were not added to ClinicalTrials.gov until November 2017 concerning FDA regulations (*Studies an FDA-regulated Device Product, Studies an FDA-regulated drug product, Device or Product Not Approved/Cleared by FDA*), and two fields that represent administrative data present in all records (*Unique Protocol Identification Number, Record Verification Date*).

Some required data element definitions were updated by the Final Rule, an amendment to FDAAA801 released on September 09, 2016. The new element definitions are legally required for all trials with start dates on or after January 18, 2017, and ClinicalTrials.gov released updated element definitions on January 11, 2017 to support the Final Rule regulations. We therefore count the number of records missing each of the 28 fields for the set of interventional records whose listed start date is before January 18, 2017, and for the set of interventional records whose start date is on or after January 18, 2017. We also divided records based on the *agency class* of the trial’s lead sponsor, which may be “NIH”, “U.S. Fed” (U.S. governmental agencies other than NIH), “Industry”, or “Other”, and counted the number of records missing required fields in each of these four categories.

We also counted the number of records with no listed “Principal Investigator”, required by the WHO dataset (called *Contact for Scientific Inquiries*), but not required by FDAAA801. A principal investigator may be listed within a ClinicalTrials.gov record either in the *responsible party* element when the *responsible party type* is “Principal Investigator” or “Sponsor-Investigator”, or in the *overall official* element.

### Eligibility Criteria

The expected format for eligibility criteria in ClinicalTrials.gov is a bulleted list of strings that enumerate the criteria below the headers ‘Inclusion Criteria’ and ‘Exclusion Criteria’. We used regular expressions to categorize the eligibility criteria from every trial record (n=302,091) as: 1) correctly formatted; 2) correct headers but not a bulleted list of criteria; or 3) missing or malformed headers (and, possibly, not formatted as a bulleted list). Out of the 117,906 records in group 2 and group 3, we manually reviewed a convenience sample of 400 records, selected at random, allowing us to extrapolate (with 95 +/-5% confidence) the number of eligibility definitions that failed to parse because they listed criteria for more than one sub-group of participants (e.g., different criteria for subjects with the studied condition and for healthy participants, different criteria for participants assigned to surgical and non-surgical intervention arms), which is not permitted in the current format.

## Results

### Simple Type Expectations

The ClinicalTrials.gov XSD schema contained type definitions for all Boolean, integer, date, and age fields, and all records validated against this XSD (**Table 3**). Therefore, all records contain correctly typed values for all occurrences of these elements. The number of fields, field names, and expected format for each simple type are displayed in **Table 3**. Dates are permitted to be “Unknown”, or similar to “January 3, 2004” or “March, 2005” where the day number is optional. Special care should be taken when parsing these dates because the behavior of many date parsing libraries when given input without a day number is undefined.

**Table 3.**
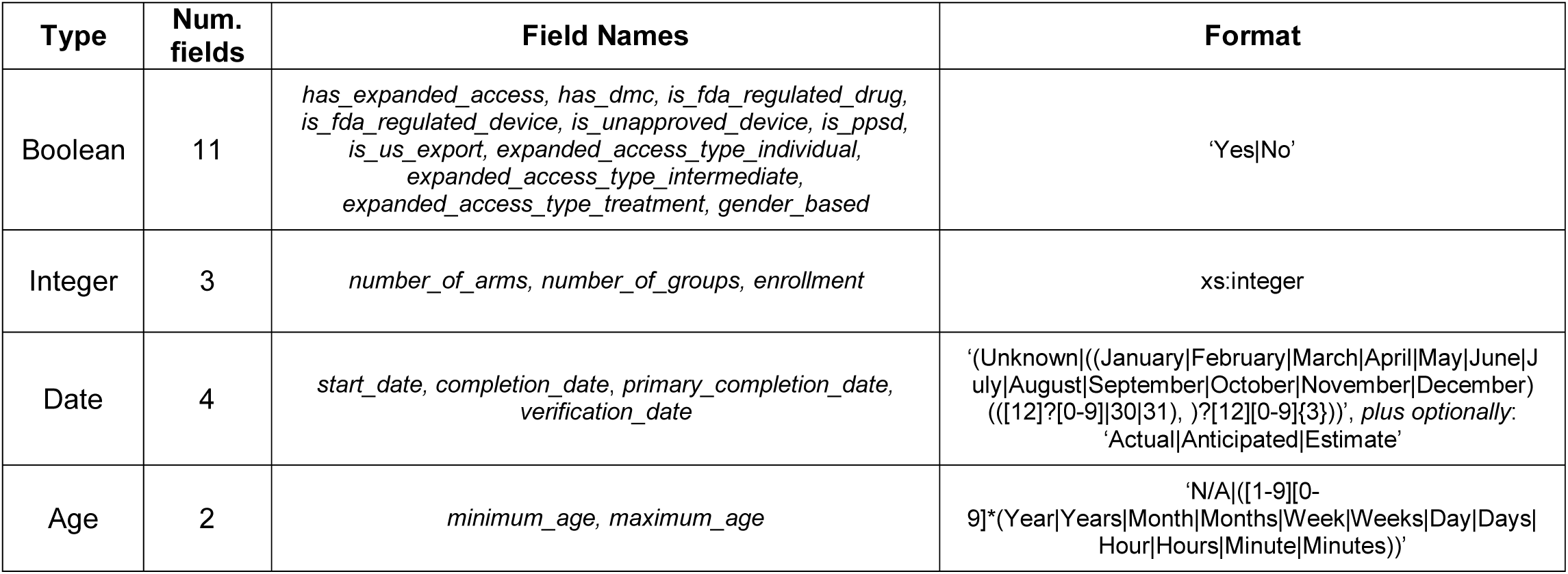
Adherence to type expectations for Boolean, integer, date, and age fields. All Boolean, integer, date, and age fields are typed in the XSD, and all values for these fields in all public records are correctly typed. Date XML elements may optionally have an attibute designating them ‘Actual’, ‘Anticipated’, or ‘Estimate’. For age fields, records may represent equivalent ages with different units (e.g., ‘2 Years’ and ‘24 Months’).

### Fields with Enumerated Values

The trial metadata contained very few “rogue” values (not drawn from the data dictionary) for fields with enumerated values (**Table 4**). Only nine of fifteen fields are typed within the XSD, however, and the untyped fields appear as free text to programs ingesting the raw XML files. For two fields, *allocation* and *masking*, the enumerated permissible values in the data dictionary use different syntax than the values that appear in the XML records (**Table 4**). The dictionary lists the acceptable values for *allocation* as “Single Group”, “Parallel”, “Crossover”, “Factorial”, and “Sequential”, but values appear in the records as “Single Group *Assignment*”, “Parallel Group *Assignment*”, etc. For the *masking* field, the data dictionary instructs the user to select from “Participant”, “Care Provider”, “Investigator”, “Outcomes Assessor”, or “No Masking”, but values appear in the actual metadata with the additional text “Single”, “Double”, “Triple”, or “Quadruple” to indicate the number of roles for people involved in the trial who are masked (or “blinded”) from knowing who has received the experimental intervention.

**Table 4.** Enumerated Value Fields Validation Results. Sixteen fields have an enumerated set of permissible values in the ClinicalTrials.gov data dictionary, but 6 are not typed within the XSD. Four of these 6 contain rogue values. For the *interventional study model* and *masking* fields, all values in public records are valid, but the data dictionary does not correctly describe the format of values. The actual format of *interventional study model* values include the word ‘assignment’ (e.g., ‘Parallel Group Assignment’ rather than ‘Parallel Group’). Values for masking include the word ‘single/double/triple/quadruple’ in addition to the types of individuals providing masking.

| Field | Valid Value Set (data dictionary) | Records With Rogue Values | Observed Rogue Values | Value Set Defined in XSD? |
| --- | --- | --- | --- | --- |
| Study Type | Interventional, Observational, Observational [Patient Registry], Expanded Access | 0 | -- | Yes |
| Overall Recruitment Status | Not yet recruiting, Recruiting, Enrolling by invitation, "Active, not recruiting", Completed, Suspended, Terminated, Withdrawn | 0 | -- | Yes |
| Responsible Party, by Official Title | Sponsor, Principal Investigator, Sponsor-Investigator | 0 | -- | Yes |
| Study Phase | N/A, Early Phase 1, Phase 1, Phase 1/Phase 2, Phase 2/Phase 3, Phase 3, Phase 4 | 0 | -- | Yes |
| Intervention Type | Drug, Device, Biologic/Vaccine, Procedure/Surgery, Radiation, Behavioral, Genetic, Dietary Supplement, Combination Product, Diagnostic Test, Other | 0 | -- | Yes |
| Sex | All, Male, Female | 0 | -- | Yes |
| Sampling Method | Probability Sample, Non-Probability Sample | 0 | -- | Yes |
| Overall Study Official's Role | Study Chair, Study Director, Study Principal Investigator | 0 | -- | Yes |
| Individual Site Status | Not yet recruiting, Recruiting, Enrolling by invitation, "Active, not recruiting", Completed, Suspended, Terminated, Withdrawn | 0 | -- | Yes |
| Interventional Study Model | Single Group, Parallel, Crossover, Factorial, Sequential | 0* | Notes: Values appear in XML as "Single Group Assignment", "Parallel Group Assignment", etc. | No |
| Masking | *Select all that apply:* Participant, Care Provider, Investigator, Outcomes Assessor, No Masking | 0* | Notes: Values appear in XML as "Double(Participant, Care Provider)", "Single(Investigator)", etc. | No |
| Primary Purpose | Treatment, Prevention, Diagnostic, Supportive Care, Screening, Health Services Research, Basic Science, Device Feasibility, Other | 196 | Educational/Counseling/Training | No |
| Allocation | N/A, Randomized, Nonrandomized | 78 | Random Sample | No |
| Arm Type | Experimental, Active Comparator, Placebo Comparator, Sham Comparator, No Intervention, Other | 21 | Case, Control, Treatment Comparison | No |
| Observational Study Model | Cohort, Case-Control, Case-Only, Case-Crossover, Ecologic or Community Studies, Family-Based, Other | 5343 | Case Control, Defined Population, Natural History | No |
| Time Perspective | Retrospective, Prospective, Cross-sectional, Other | 622 | Longitudinal, Retrospective/Prospective | No |

### Completeness

We counted the number of records missing each field for 28 of the 41 fields required by the FDAAA801 Final Rule. Fourteen of these required fields (*Brief Title, Brief Summary, Study Phase, Study Type, Primary Disease or Condition Being Studied, Intervention Name(s), Intervention Type, Eligibility Criteria, Sex/Gender, Age Limits, Overall Recruitment Status, Name of the Sponsor, Responsible Party by Official Title, Secondary ID*) are present in all records or missing in a negligible number of records (<.03%). The numbers of records missing the remaining fourteen required fields (missing in a non-negligible number of records) are displayed in **Table 5**.

**Table 5.** Missing required fields, before and after passage of FDA Final Rule. For fields required by the FDAAA801 Final Rule, table lists the percentage of all interventional records (n=239,274) missing the field, and the percentage of all interventional records with start dates after the effective date of the Final Rule (n=46,289) missing the field. **m** indicates multiple instances of field are permitted; a multiple field is considered ‘missing’ if there are no listed occurrences of field. **c** indicates a conditionally required element, such as *Why Study Stopped*, which is required only if the study terminated before its expected completion date. Conditionally required elements are considered missing if they are both missing and conditionally required for the given record.

| Required Field Name |  | Number of Interventional Records Missing Field |  | Percentage of Records Missing Field |
| --- | --- | --- | --- | --- |
|  |  | Trials starting before 01/18/17, effective date of Final Rule (n=192985) | Trials starting on or after 01/18/17 (n=46289) | All Interventional Trials (n=239274) |
| **(i) Descriptive Information** |  |  |  |  |
| (B) Official Title |  | 7038 | 10 | 2.9% |
| (D) Primary Purpose |  | 8253 | 6 | 3.5% |
| (E) Study Design | interventional study model | 6834 | 0 | 2.9% |
|  | number of arms | 23,832 | 296 | 10% |
|  | allocation | 44,552 | 11938 | 24% |
|  | masking | 5423 | 1 | 2.3% |
|  | arm information **m** | 23,832 | 296 | 10% |
| (L) Intervention Description, for each intervention studied **m** |  | 45,433 | 29 | 19% |
| (S) Study Start Date |  | 2950 | 3 | 1.3% |
| (T) Primary Completion Date |  | 15,044 | 0 | 6.3% |
| (U) Study Completion Date |  | 13,516 | 21 | 5.7% |
| (V) Enrollment |  | 3966 | 0 | 1.7% |
| (W) Primary Outcome Measure Information **m** | outcome measures *missing* | 7,786 | 0 | 3.3% |
|  | time frame | 12,865 | 0 | 5.4% |
|  | description | 82,862 | 5,086 | 37% |
| **(ii) Recruitment Information** |  |  |  |  |
| (D) Accepts Healthy Volunteers |  | 1,323 | 0 | .55% |
| (F) Why Study Stopped **c** |  | 4,385 | 2 | 1.8% |
| (G) Individual Site Status **c m** |  | 0 | 0 | 0% |
| (H) Availability of Expanded Access |  | 3,158 | 833 | 1.7% |
| **(iii) Location and contact information** |  |  |  |  |
| (C) Facility Information **m** | no listed facilities | 20,351 | 5,602 | 11% |
|  | facility name, city, or country *missing* | 7,899 | 17 | 3.3% |
|  | Overall contact info OR contact info for each site | 163,905 | 13,123 | 74% |

Several elements were added or updated in the Final Rule. *Official title, why study stopped* (for a study that is “suspended”, “terminated” or “withdrawn” prior to its planned completion), *study start date*, and *study completion date* were recommended but not required for studies with start dates before the effective date of the Final Rule. FDAAA801 defines a single study design element whose textual definition states that study design information should include “Interventional study model, number of arms, arm information, allocation, masking, and whether a single-arm trial is controlled.” This information was stored in ClinicalTrials.gov records as a comma-separated list. The Final Rule eliminates the single-arm control element, makes *interventional model, allocation, enrollment*, and *masking* required sub-elements of *study design*, and makes *number of arms* and *arm information* separate required elements.

The curators at the NLM have parsed the study design information in historical records and inserted it into the new structured element. However, there is no way to confirm that this conversion was done with complete accuracy, and any unstructured data in the original study design elements have been lost, even within the historical versions of records that ClincalTrials.gov provides. *Arm information* consists of a *label* (e.g., the name of the experimental intervention used or placebo), *type* (“Experimental”, “active comparator”, “placebo comparator”, or “other”), and *description. Label* and *type* were present in all listed arms, but 10% of the records failed to list any study arms.

Most required fields are present in nearly all records submitted after the January 2017 update to ClinicalTrials.gov because automated PRS validation rules prevent the submission of records with missing required fields. However, contact information was frequently missing or underspecified both before and after the Final Rule. Contact information consisting of a *name, phone number*, and *email address* is required either for each individual facility, or a single *overall contact* is required for the trial. No *overall contact* is given in 223,468 trials (74% of all trials), and of those trials only 5,475 list a primary contact for each trial location. Of all 385,279 contact details that are provided, either as the *overall contact* or a location-specific contact, 81,195 (21%) lack a phone number and 86,611 (22%) lack an email address.

We also counted the number of interventional trial records missing values from each of the 41 fields required by the Final Rule after categorizing the records based on the *agency class* of the lead sponsor. We found 6,851 trials sponsored by the NIH, 3,032 trials sponsored by “U.S. Fed” (US governmental agency other than NIH), 69,100 trials sponsored by industry, and 160,291 trials with *agency class* “other”. We found that records from trials with a lead sponsor of “NIH” contain significantly more missing values than do those from the other three agency classes. The fields for which the difference in the number of records missing a value for the field across agency class is the greatest are displayed in **Figure *2***. There was not a significant difference in completeness of other fields across agency class of lead sponsor; record counts for all missing fields for all agency classes are included in the supplementary material.

**Figure 2.**
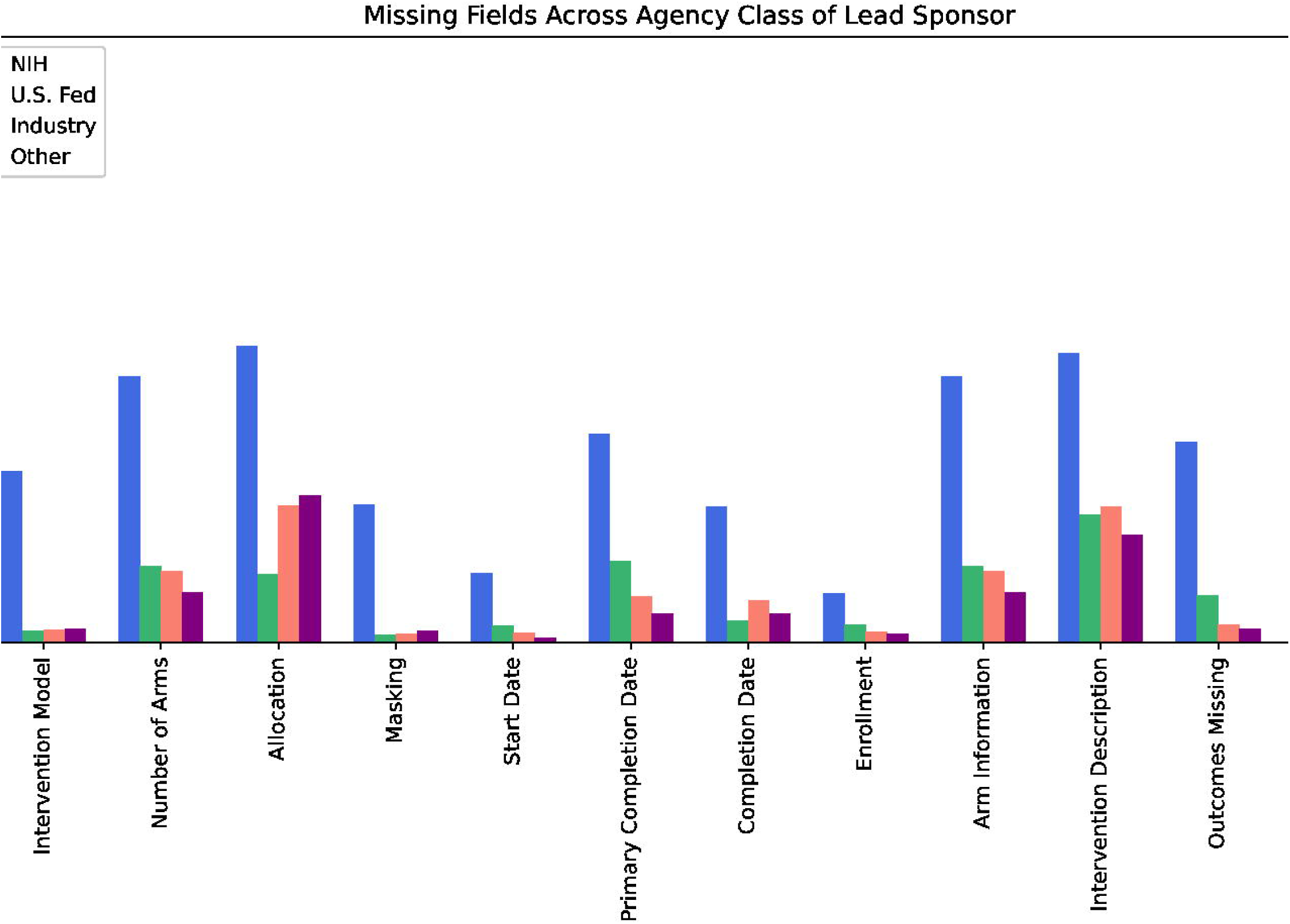
Percentage of Interventional Records Missing Required Field Values, by *Agency Class* of Lead Sponsor. Percentage of ClinicalTrials.gov interventional trial records (n=239,274) missing values for selected fields required by FDAAA801. Records are categorized by the *agency class* of the lead sponsor, which is either “NIH”, “U.S. Fed”, “Industry”, or “Other”.

The eighth element in the WHO dataset is a *Contact for Scientific Queries*, and its description states that there must be “clearly assigned responsibility for scientific leadership to a named Principal Investigator” and that this data element must include “Name and title, email address, telephone number, postal address and affiliation” of the PI, even if additional contact details are provided for a second contact. The FDA requires only a responsible party, which is permitted to be a sponsor. If the *responsible party type* is “Principal Investigator” or “Sponsor-Investigator”, then only *name, investigator affiliation*, and *investigator title* are required. ClinicalTrials.gov provides an additional element, *overall official*, which corresponds to the WHO PI element, but it is not required, and the element definition does not include contact information.

Out of 302,091 trials, 35,226 (12%) have no listed principal investigators, 22,557 (7.5%) have a *responsible party type* of “Principal Investigator” or “Sponsor-Investigator” but do not separately designate an *overall official*, 162,985 (54%) have an *overall official* and a non-scientific responsible party (e.g., a sponsor), and 81,323 (27%) list both an *overall official* and *investigator information* for the responsible party.

We noticed irregularities in the structure of both investigator and contact-related elements. Within the XSD, an investigator has sub-fields *first name, middle name, last name, degrees, role, affiliation*, and a contact has sub-fields *first name, middle name, last name, degrees, phone, phone ext*, and *email*. However, *first name, middle name*, and *degrees* are missing in all investigators and contacts in all records, and instead the individual’s full name and degrees all appear within the value of the *last name* field (e.g., “Sarah Smith, M.D.”). The *responsible party* element contains sub-fields *investigator affiliation, investigator full name*, and *investigator title* with no field for degrees.

### Ontology-Restricted Fields

The only field currently restricted to terms from an ontology is the *condition* field. Rather than being restricted to a single ontology, authors are encouraged to use either MeSH terms, or terms than can be mapped to MeSH through the UMLS metathesaurus. Within the ClinicalTrials.gov records, values for *condition* appear as simple strings (e.g., “diabetes mellitus”) rather than as globally unique, persistent identifiers. During metadata creation, PRS attempts to map user-submitted *condition* strings to UMLS concepts. If the mapping is successful, PRS accepts the user string as-is, without including the UMLS concept identifier in the metadata or replacing the user string with a standard syntactic representation of the concept. Alternative spellings (“tumor” vs “tumour”) and synonyms (“breast cancer” vs “malignant neoplasm of the breast”) are not harmonized.

ClinicalTrials.gov addresses searchability issues that would normally arise in a database containing unharmonized synonyms by building a computation engine into its search portal that parses queries for UMLS concepts, and includes synonyms in the search (**Table 6**). While this system mitigates some of the issues arising from synonyms and alternative spellings, it is available only through the ClinicalTrials.gov search portal, and unharmonized values persist in the raw metadata. Only synonyms for the query term are provided, and users cannot browse from their original query to more or less general concepts (e.g., in the MeSH hierarchy) in order to refine their search. These detected synonyms are always included and the user cannot choose to search for an exact phrase.

**Table 6.**
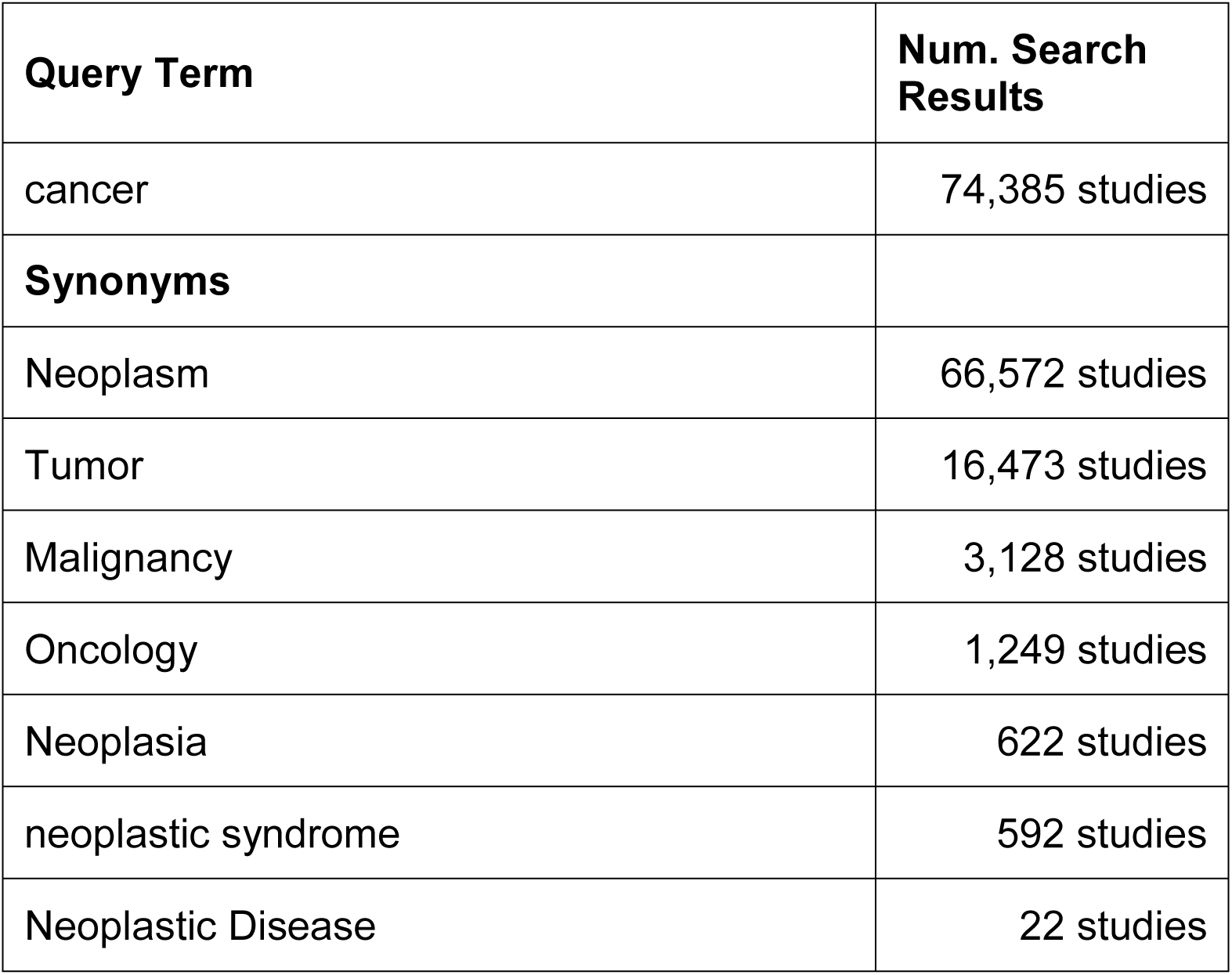
Synonyms Added by ClinicalTrials.gov for Search Term “Cancer”. Behind the scenes, the ClinicalTrials.gov search portal adds 7 synonyms to a user query for “cancer”. Note: Inconsistent capitalization accurately reflects how terms are displayed in ClinicalTrials.gov.

We performed a comprehensive search for concepts in UMLS ontologies that matched the values given for the *condition* field. First, we checked adherence to the restrictions as they are defined. We found that only 306,197 (62%) of the 497,124 listed conditions return an exact match in MeSH, and only 402,875 (81%) have an exact match in any UMLS-mapped ontology. Second, we evaluated whether any UMLS ontology alone was sufficient to provide all terms used for the condition field (**Figure 3**). Of the 190,927 condition terms that have no match in MeSH, 96,678 conditions (51%) do have an exact match in another ontology. MeSH provides the best coverage of any single ontology, but it does not cover significantly more terms than MEDDRA, which contains matches for 230,639 conditions (46%), or SNOMED-CT, which contains matches for 224,008 conditions (45%).

**Figure 3.**
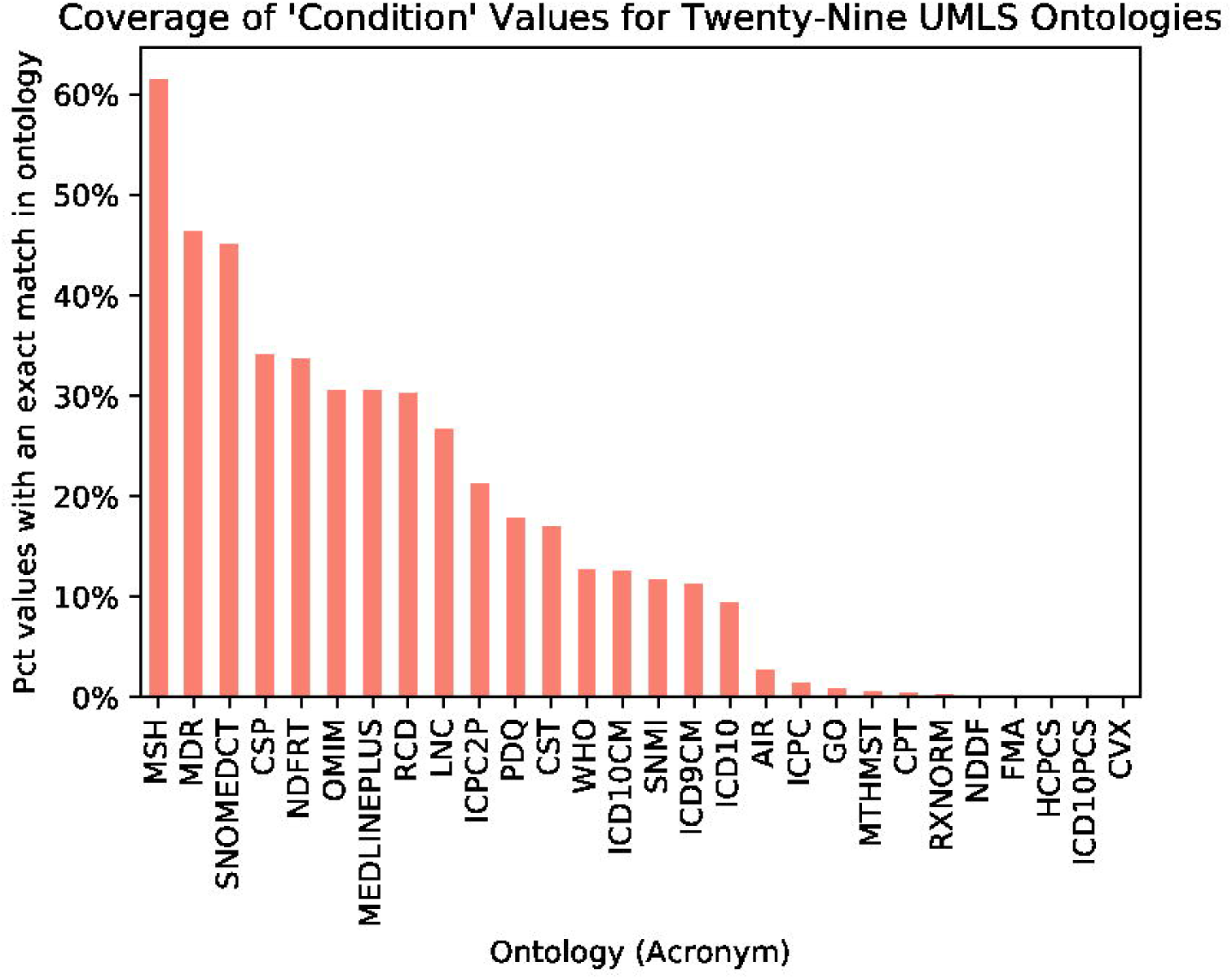
Percentage of values for the *condition* field covered by each UMLS Ontology. Each column gives the percentage of the 497,124 values for the *condition* field contained in ClinicalTrials.gov records that are an exact match for a term from the given ontology. Of 72 ontologies in the UMLS, 29 contained at least one match for a *condition* value, and 43 contained no matches (omitted from figure). Many *condition* values have exact matches in more than one ontology. The ontologies that provide the most coverage for *condition* values are MeSH (62%), MedDRA (46%), and SNOMED-CT (45%).

We verified that the *intervention* field could be restricted to ontology terms without significant loss of specificity, by demonstrating that 256,463 out of 557,436 listed intervention values (46%) can be matched to terms from BioPortal ontologies, even without any pre-processing (**Figure 4**). All interventions have an associated *intervention type*, one of the eleven choices in **Figure 4**, and usage of ontology terms varies greatly between types.

**Figure 4.**
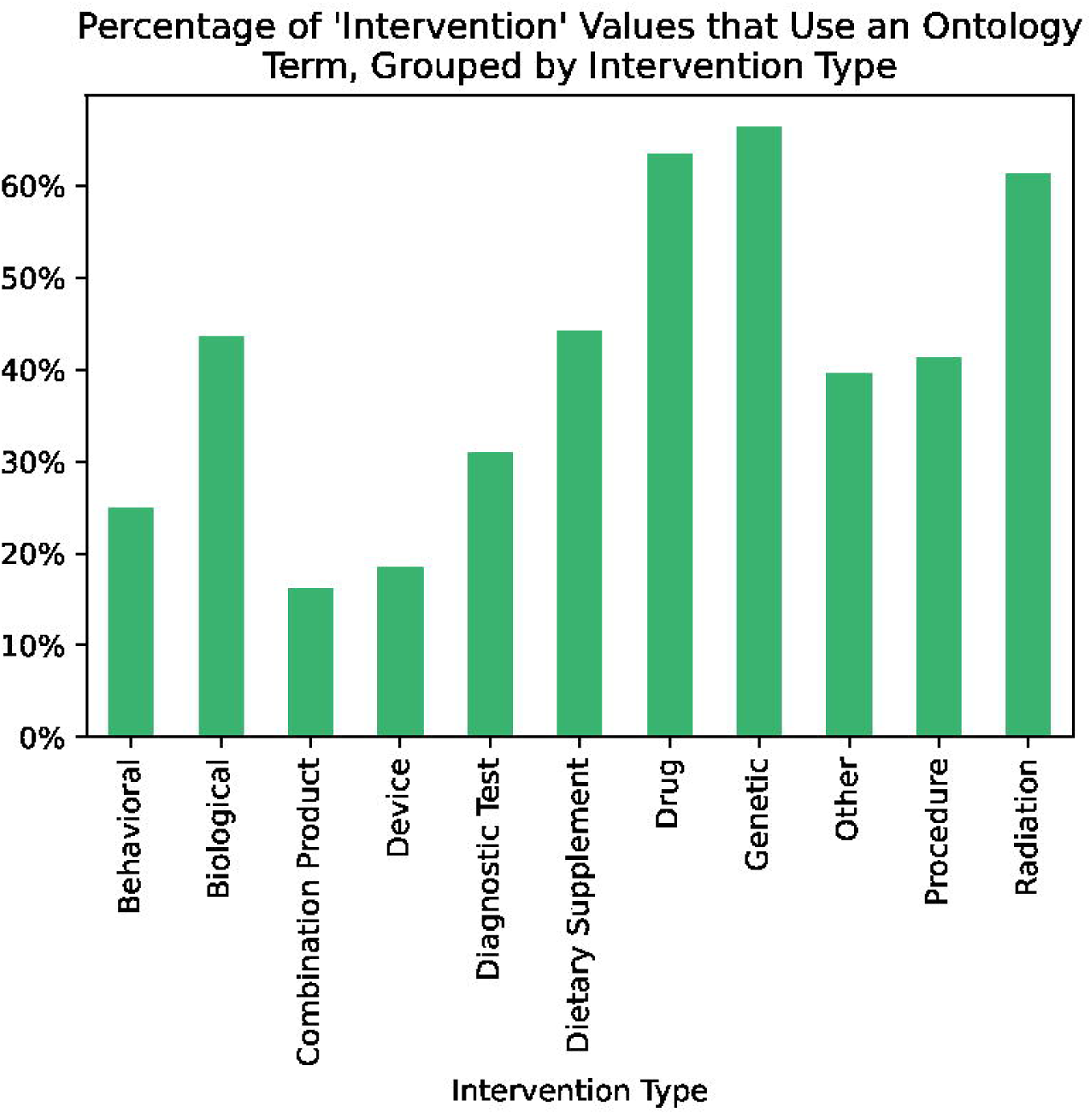
Percentage of values for the *interventio*n field for which we found an exact match in at least one ontology hosted in NCBO BioPortal, grouped by intervention type. Thirty-nine percent of the 557,436 values listed for *intervention* contained in ClinicalTrials.gov records are an exact match to a term from a BioPortal ontology, without any pre-parsing or normalization, indicating that this field could reasonably support ontology restrictions. Some intervention types are much better represented by ontology terms than others. More than half of all drugs and radiation therapies use ontology terms, but less than 15% of listed devices and combination products do.

### Eligibility Criteria

The eligibility-criteria element is a block of semi-formatted free text. The data dictionary says to “use a bulleted list for each criterion below the headers ‘Inclusion Criteria’ and ‘Exclusion Criteria’”, and the PRS prepopulates a textbox with the correct format. However, there is no enforced format restriction. We find that only 183,312 records (61%) follow the expected format for eligibility criteria (**Table 7**). Error types include:

**Table 7.** Number of records with missing and incorrectly formatted eligibility criteria. Table shows the count and percentage of ClinicalTrials.gov records with correctly formatted criteria, missing criteria, and the two most common incorrect formats: incorrect ‘inclusion’ and ‘exclusion’ headers, and non-bulleted criteria.

| Number of Records (n=302,091) |  |  |  |
| --- | --- | --- | --- |
| Correctly Formatted Eligibility Criteria | Correct headers, but not formatted as a bulleted list | Missing or Malformed Headers | Missing Eligibility Criteria |
| 183,309 (60.7%) | 73,771 (24.4%) | 44,135 (14.6%) | 876 (.29%) |

- Missing one or both inclusion/exclusion headers
- Misspelled or alternative inclusion/exclusion headers
- Criteria not formatted, or only partially formatted as a bulleted list
- Criteria defined for sub-groups of participants, and/or defined for non-subjects

We manually reviewed a random sample of 400 of the 117,906 records from the second and third groups in **Table 7** (i.e., all records in which eligibility criteria were present but incorrectly formatted). Of this sample, 55 records (14%) defined separate criteria for sub-groups of participants (e.g., subjects with the studied condition and healthy participants, participants assigned to surgical arms and participants assigned to non-surgical arms). Based on these results, we estimate that 16,200 records, 5% of records in the entire repository, define eligibility criteria for multiple groups of participants. The most common cause of criteria failing to parse according to the expected format was paragraph-style sentences interspersed with bulleted criteria.

## Discussion

The metadata in ClinicalTrials.gov are of higher quality than the metadata in other biomedical repositories that we have examined^42^. Values for numeric, date, and Boolean fields, and fields with enumerated permissible values conform to type expectations. The presence of some required fields is strictly enforced by the Protocol Registration System, but contact information, principal investigators, study design information, and outcome measures are frequently missing. Two key fields for searching records, *condition* and *intervention*, do not restrict values to terms from ontologies, so users cannot easily refine or broaden search queries. Eligibility criteria are stored as semi-structured text; they are recommended to be formatted as a bulleted list of individual criteria, but nearly 49% of values fail to parse according to the expected format.

In NCBI’s BioSample repository and Gene Expression Omnibus (GEO), and in EBI’s BioSamples repository, the two issues most impeding data reuse are non-standardized field names and malformed values that failed to conform to the expected type for their field^41,42^. Apart from minor irregularities in some fields with enumerated values, ClinicalTrials.gov metadata were entirely free from these issues. In all cases, the design of the metadata authoring system played a key role in the quality of the metadata. BioSample, BioSamples, and GEO all provide templates suggesting a particular format, but they do not enforce restrictions, placing that burden on metadata authors. In contrast, the PRS provides automated validation for most fields, immediately displays error messages to metadata authors, and does not allow records with outstanding errors to be submitted. Other metadata repositories would benefit from using similar techniques in their data-entry pipelines. The Center for Expanded Data Annotation and Retrieval (CEDAR)^59^ has created such a platform for metadata authoring—similar to PRS in that it enforces a schema and is based on forms. The advantage of CEDAR is that it provides tight integration with biomedical ontologies to control both field names and values, while not being tied to a single repository or metadata schema.

Restricting the allowed values for certain fields in biomedical metadata to terms drawn from an ontology, such as MeSH, can improve metadata computability and reusability by eliminating the usage of synonyms and preventing typographical errors, providing a defined range of values over which analyses can be performed, and enabling users to develop and refine search queries by navigating up or down in the term hierarchy. No field within ClincialTrials.gov records is required to use values from an ontology, and the data dictionary recommendation that the *condition* field use values that “can be mapped to MeSH” through the UMLS Metathesaurus is too vague to provide a defined set of expected values. Even when values for fields in ClinicalTrials.gov records are drawn from an ontology, they are not specified using globally unique and persistent identifiers, which would enable the interoperability of data with systems that expect these well-defined terms as input. Our results demonstrate that there is no single ontology that covers the majority of needed terms (**Figure 3**). One possible solution would be to define a custom extension to MeSH that adds additional values to the existing hierarchy that can be used to populate the *condition* field.

For both the *condition* and *intervention* fields, ClinicalTrials.gov addresses searchability issues caused by the existence of synonyms in field values, and the lack of a defined range of search terms, by automatically rewriting queries to include synonyms of the terms provided by the user (**Table 6**). However, this functionality only exists in the ClinicalTrials.gov search portal. The synonyms are not included in the raw XML records obtained from ClinicalTrials.gov. Consequently, search results using the raw trial records as opposed to the ClinicalTrials.gov portal can be radically different. Moreover, the inclusion of synonyms in searches cannot be toggled off, and users may disagree with ClinicalTrials.gov’s definition of synonymy. For instance, it is debatable whether a “tumor”, which may be benign, is a synonym for “cancer”, of which not all types result in tumors. “Tumor” is always included in user queries for “cancer”, however (**Table 6**).

In December, 2019 the National Library of Medicine issued a Request for Information (RFI) to obtain suggestions for modernizing ClinicalTrials.gov. Results from this RFI were made publicly available in April, 2020^60,61^. As we do, several respondents suggested standardizing the vocabulary used in records by encouraging greater use of well-known controlled terminologies. Respondents also requested the ability to search for an exact phrase (i.e., without synonyms) and the ability to search by disease subtype (which restricting values to a hierarchy such as MeSH would provide).

We found that the reusability of clinical-trial metadata is hindered by the lack of a single minimum information standard for the fields required for registering clinical trials, and by the discrepancies between the 24 fields required by the WHO Trial Registration Data Set^52^ and the 41 fields required by FDAAA801. FDAAA801 does not require a principal investigator or contact for scientific inquiries to be listed, and 12% of interventional trials in ClinicalTrials.gov fail to list any principal investigator. Many of the fields that are shared between the WHO data set and FDAAA801 have different names and definitions within the two standards. Both standards have had multiple updates and changes to required elements since their first publication, so that older records in *ClinicalTrials*.*gov* (especially those added before the Final Rule-related site update) are often missing fields that were added or became required later. The maintainers of ClinicalTrials.gov have done an admirable job of parsing unstructured data such as the study design information from old records into the new structured format, but details from the original unstructured data have almost certainly been lost. Automated validation rules in the PRS system have been moderately successful at ensuring required fields are filled for trials after January 18, 2017, but important fields such as the method of *allocation* of patients to study arms, and contact information are still often missing (**Table 5**).

Metanalyses involving the principal investigators of trials are further hindered by the fact that principal investigator information may be listed as part of either the *responsible party* field, the *overall official* field, both, or neither. Further, *responsible party* uses a single sub-field (*investigator full name*) to store the entire name, but *overall official* has sub-fields *first name, middle name*, and *last name*. Like Chaturvedi et al.^47^, we recommend that investigator information be augmented or replaced with persistent unique identifiers such as Open Researcher and Contributor IDs (ORCID). Using ORCIDs would ensure that investigator information is consistent across multiple trials and across multiple listings in the same record. Currently within the PRS, investigator information must be manually reentered everywhere it occurs, which may validly be as the *responsible party*, as the *overall official*, and as the site-specific investigator for one or more locations for a single trial. Using ORCIDs also would allow for a researcher’s name, degrees, and affiliation to change with time and to be simultaneously updated in all records.

Similarly, fields that may refer to a research organization (*sponsors, collaborators, investigator affiliation* for the *responsible party, affiliation* for the *overall official*, and *facility name* for trial locations) could be augmented with identifiers from the Research Organization Registry (ROR). The ROR is a community-led project to develop an open, sustainable, usable, and unique identifier for every research organization in the world, which provides mappings to other identifiers (e.g., GRID, ISNI, Wikidata), and a publicly accessible API for querying the registry^62^. However, adding either ORCIDs or RORs to ClinicalTrials.gov records will present challenges such as disambiguating values in existing records (e.g., which of several ORCIDs for “John Smith” is correct?) and determining whether ORCID and ROR entries should be created for entities if they do not exist.

Some respondents to the NLM’s RFI requested the ability to search for studies by eligibility criteria, a more prominent and detailed display of eligibility criteria, or structured information for the inclusion and exclusion criteria. Developing structured representations of inclusion and exclusion criteria that may be reused in future studies, or used to automatically match eligible patients (e.g., from a hospital’s patient database) is an active area of research^63,64^. Cohort definition and recruitment are among the most challenging aspects of conducting clinical trials,^65^ and difficulties in recruitment cause delays for the majority of trials^66,67^. Several groups have developed structured representations and grammars for eligibility criteria^64,68^, all of which require that criteria are structured as combinations of Boolean criteria, but there is no expert consensus on a representation system, nor is it reasonable to expect the principle investigators of most trials to formulate criteria in this logically precise way.

There are, however, several improvements that could be made to the *eligibility criteria* field in ClinicalTrials.gov that would facilitate searching records by this field, and using natural language processing methods to extract structured Boolean criteria from the semi-structured text at a later date, First, *inclusion criteria* and *exclusion criteria* should become separate fields. This change would eliminate the need for user-defined headers, and fix 37% of existing errors, as **Table 7** shows. Further, the PRS should allow users to enter multiple blocks of inclusion and exclusion criteria and an associated criteria group name, such as the label of the corresponding study arm. Our manual review of the *eligibility criteria* element suggested that at least 5% of trials have criteria for multiple groups. However, our analyses assumed that records with correctly formatted eligibility criteria never required multiple sets of criteria. There are almost certainly trials in which the full protocol specified eligibility criteria for multiple groups, but the metadata author entered a simplified version of the protocol due to the lack of an appropriate field, so the true percentage is likely much higher.

Our work is limited to the clinical-trial protocols stored in ClinicalTrials.gov, and could be expanded to cover adherence to schema, missing fields, and usage of ontology terms in the summary results of ClincialTrials.gov records, which are required to be submitted within one year of the study completion date. Since we are primarily concerned with the reusability of the existing metadata, we did not evaluate whether protocol elements and results were added in a timely manner in accordance with FDAAA801 and the Final Rule. Several past studies have shown low levels of compliance with mandatory results reporting^11,44,69^, and DeVito et al.^70^ found that, in data taken from ClinicalTrials.gov on September, 2019, only 1722 out of 4209 applicable trials due to report results had reported results by the 1-year deadline. Both DeVito and Anderson^69^ found that lower levels of compliance with results reporting were associated with trials funded by NIH and other US governmental institutions versus trials funded by industry. This is consistent with our findings that trials with a lead sponsor within the NIH were more likely to be missing required fields (**Figure 2**).

Although the registrations in ClinicalTrials.gov serve the important purpose of enabling FDA oversight and protecting human subjects, they are also an invaluable source of metadata about clinical trials for systematic reviews, adverse events^10,11^, and analyses about funding sources^12–16^, study design^17,19,22^, time to publication following study completion^11,23–26^, geographical availability of trials^27–32^, causes of delays and early terminations^33–40^, and more. ClinicalTrials.gov is also an important resource for patients and health care providers to search for studies for which patients are eligible. It is therefore encouraging that the NLM has stated their intention to modernize ClinicalTrials.gov, with a focus on improving data interoperability and reuse, and on serving needs and users beyond its original purpose. Our analysis highlights the limitations of the current metadata stored in ClinicalTrials.gov and the benefits that would ensue from making ClinicalTrials.gov records more structured, and thus more findable by specific searches, interoperable with other knowledge sources, and reusable in statistical analyses of multiple studies.

## Code Availability

Software used to query the BioPortal API extends an existing suite of metadata analysis tools maintained by the Center for Expanded Data Annotation and Retrieval (CEDAR) and is available at https://github.com/metadatacenter/metadata-analysis-tools/. Python notebooks which reproduce all other analyses, tables, and figures are available at https://github.com/lauramiron/CTMetadataAnalysis.

## Data Availability

The data used and generated throughout the study described in this paper are available in Figshare at <u>10.6084/m9.figshare.12743939</u>^71^. Stanford University’s PRS test environment was used for qualitative analysis of the PRS system. This environment is not publicly accessible, but our results can be verified in any PRS environment to which the reader has access.

## Acknowledgments

This work was supported by American Heart Association Institute for Precision Cardiovascular Medicine grant 18IDHP34660267, by the U.S. National Institutes of Health under grants AI117925 and GM21724, and in part by Stanford CTSA Award UL1TR003142 from the National Center for Advancing Translational Science (NCATS), a component of the National Institutes of Health.

## Author Contributions

L.M.: Study design, data collection, software implementation, study execution, data analysis, and manuscript writing. R.S.G.: Study design, software implementation, manuscript review and editing. M.M.: Conceptualization, study design, manuscript review and editing.

